# ‘Nepotistic journals’: a survey of biomedical journals

**DOI:** 10.1101/2021.02.03.429520

**Authors:** Alexandre Scanff, Florian Naudet, Ioana Cristea, David Moher, Dorothy V M Bishop, Clara Locher

## Abstract

**Context:** Convergent analyses in different disciplines support the use of the Percentage of Papers by the Most Prolific author (PPMP) as a red flag to identify journals that can be suspected of questionable editorial practices. We examined whether this index, complemented by the Gini index, could be useful for identifying cases of potential editorial bias, using a large sample of biomedical journals.

**Methods:** We extracted metadata for all biomedical journals referenced in the National Library of Medicine, with any attributed Broad Subject Terms, and at least 50 authored (i.e. by at least one author) articles between 2015 and 2019, identifying the most prolific author (i.e. the person who signed the most papers in each particular journal). We calculated the PPMP and the 2015-2019 Gini index for the distribution of articles across authors. When the relevant information was reported, we also computed the median publication lag (time between submission and acceptance) for articles authored by any of the most prolific authors and that for articles not authored by prolific authors. For outlier journals, defined as a PPMP or Gini index above the 95th percentile of their respective distributions, a random sample of 100 journals was selected and described in relation to status on the editorial board for the most prolific author.

**Results:** 5 468 journals that published 4 986 335 papers between 2015 and 2019 were analysed. The PPMP 95^th^ percentile was 10.6% (median 2.9%). The Gini index 95^th^ percentile was 0.355 (median 0.183). Correlation between the two indices was 0.35 (95CI 0.33 to 0.37). Information on publication lag was available for 2 743 journals. We found that 277 journals (10.2%) had a median time lag to publication for articles by the most prolific author(s) that was shorter than 3 weeks, versus 51 (1.9%) journals with articles not authored by prolific author(s). Among the random sample of outlier journals, 98 provided information about their editorial board. Among these 98, the most prolific author was part of the editorial board in 60 cases (61%), among whom 25 (26% of the 98) were editors-in-chief.

**Discussion:** In most journals publications are distributed across a large number of authors. Our results reveal a subset of journals where a few authors, often members of the editorial board, were responsible for a disproportionate number of publications. The papers by these authors were more likely to be accepted for publication within 3 weeks of their submission. To enhance trust in their practices, journals need to be transparent about their editorial and peer review practices.

## Introduction

In the field of academic publishing, the term ‘self-promotion journal’ was coined to reflect the dubious editorial practices of New Microbes and New Infection (NMNI), an Elsevier journal, whose most prolific author, Didier Raoult, co-authored 32% of the 728 published papers. NMNI’s editor-in-chief and six additional associate editors of the journal work directly for Raoult. Together, they authored 44% of the 728 papers published up to June 25, 2020 in the journal. ‘Self-promotion journals’ were therefore proposed as a new type of illegitimate publishing entity, which could have certain key characteristics such as (i) a constantly high proportion of papers published by the same group of authors, (ii) relationships between the editors and these authors, and (iii) publication of low-quality research.

We applied a preliminary approach to detect ‘self-promotion journals’ in the field of infectious disease using a simple measure: the proportion of contributions published in a journal by the most prolific author, i.e. the one who published the most articles in a given time period.[1] In journals publishing more than 50 papers over 5 years, it was rare to see journals where a specific author published more than 10% of the papers, and indeed NMNI was a clear outlier. One of our team (DB) reported a similar analysis for the addiction subfield of psychology[2] in a blog post. She found a bimodal distribution of ‘the percentage by the most prolific’ measure, identical to that observed for NMNI, with only 3 out of 99 journals having a score over 8%. In two of these journals, the high score was attributable to the same individual, who was on the editorial board of the journal, and who had published together with the editor-in-chief. Bishop also noted that the same method identified a journal editor, Johnny Matson, who had been found previously to be publishing copiously in journals edited by himself or other editors[2]. Furthermore, many of these papers, with superficial or absent peer reviews, could be detected by the remarkably rapid turn-around, often within a week or less, between the dates recorded for submission and acceptance. This additional evidence of unethical editorial practice can only be obtained, however, in journals that report these dates for published manuscripts.

These convergent analyses from different fields suggest that the Percentage of Papers by the Most Prolific author (PPMP) is a simple measure that can be used as a red flag to identify journals that are suspected of biased editorial decision-making – what we now term ‘nepotistic journals’. The PPMP is also in line with studies on resource distribution in economics. A highly prolific author who is an outlier on the PPMP measure monopolizes a large part of a journal’s publications. This analogy supports another, more complex measure, the Gini index[3], used in econometrics to describe resource distribution inequalities. Applied to our context, it could be used to quantify imbalances in the patterns of authorship within a journal.

We set out to apply these two indices to a very large dataset of biomedical journals over a 5-year timeframe, to describe outliers using these indices and to describe time intervals between submission and acceptance dates as a potential surrogate for unfair or partisan editorial practices.

## Results

### Journal selection and description

Using the search query on the United States National Library of Medicine catalog, 11 665 journals labelled with at least one of 152 ‘Broad Subject Terms’ were retrieved. **Figure 1** details the reasons for non-inclusion of some journals. After exclusions, 5 468 journals were analysed.

**Fig 1:**
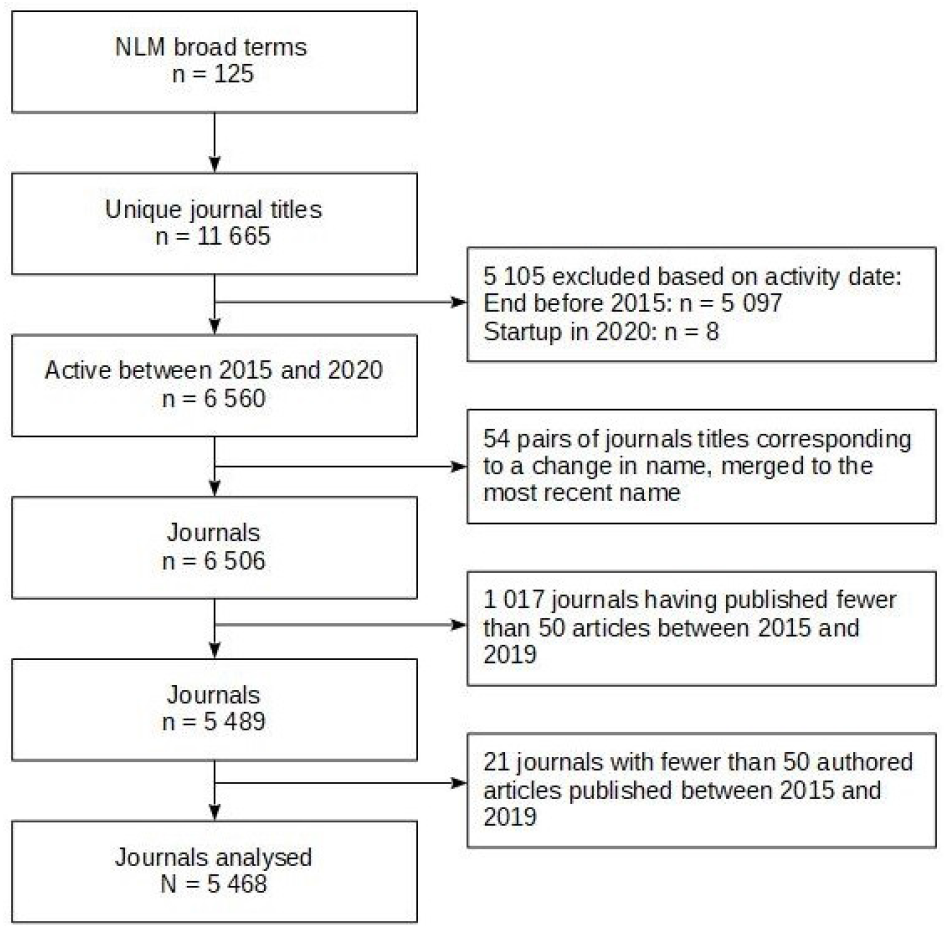
Selection flowchart for journals labelled with at least one ‘Broad Subject Term’ in the National Library of Medicine (NLM).

These journals published a total of 4 986 335 articles of which 4 582 473 were ‘journal articles’ (**see Web-Appendix 1**). The main characteristics of the journals analysed are described in **Table 1**. Briefly, they published a median of 500 articles (IQR 262 to 964) for the 2015–2019 period, of which 463 (IQR 246 to 876) were considered ‘journal articles’. Two ‘mega-journals’ published more than 25 000 articles over the 5-year period (*Scientific Reports* with 95 890 articles, and *PloS One* with 107 342 articles). For 3668 journals (67%), there was at least one article without any author, and a median percentage of 0.9% of articles (IQR 0.4% to 2.1%) with no named author in these journals. The author with the largest number of articles in a given journal is a journalist (N = 767) and the author with the largest number of ‘journal articles’ is an academic (N = 471). Both authors publish in established journals (*The BMJ* and *Journal of the American Dental Association*, respectively) and are members of their respective editorial boards.

**Table 1:** Main characteristics of all journals in the United States National Library of Medicine catalog having at least one Broad Subject term and having published at least 50 authored articles between 2015 and 2019.

|  | All articles | Journal Articles |
| --- | --- | --- |
| **Number of journals with $\geq 50$ authored articles** | 5468 | 5455 |
| **Number of articles** |  |  |
| Median [IQR] | 500 [262 - 964] | 463 [246 - 876] |
| Range | 50 - 108 990 | 50 - 105 031 |
| **Number of articles with an author** |  |  |
| Median [IQR] | 494 [257 - 952] | 458 [243 - 868] |
| Range | 50 - 107 342 | 50 - 104 713 |
| **PPMP (%)** |  |  |
| Median [IQR] | 2.88 [1.71 - 4.91] | 2.67 [1.65 - 4.48] |
| Range | 0.1 - 39.9 | 0.1 - 42.7 |
| **PPMP 95th percentile (%)** | 10.6 | 9.6 |
| **Number of articles by MPA** |  |  |
| Median [IQR] | 14 [8 - 25] | 12 [8 - 21] |
| Range | 1 - 767 | 1 - 471 |
| **Tied as MPA** | 1 022 (19%) | 1 156 (21%) |
| **Gini index** |  |  |
| Median [IQR] | 0.183 [0.131 - 0.246] | 0.172 [0.123 - 0.232] |
| Range | 0.00 - 0.74 | 0.00 - 0.72 |
| **Gini index 95th percentile** | 0.355 | 0.343 |
| **Median of publication lag ratio (MPA / no MPA)** |  |  |
| Median [IQR] | 0.829 [0.606 - 1.02] | 0.846 [0.647 - 1.03] |
| Range | 0.0 - 26.8 | 0.0 - 7.7 |
| Not calculable | 2743 (50%) | 2768 (51%) |

### Description of the Indices

#### Percentage of papers by the most prolific author and Gini index

For the 2015–2019 period, the PPMP median was 2.88% (IQR 1.71% to 4.91%) and the 95 ^th^ percentile of the PPMP value was 10.6% (**Figure 2**). The most prolific author(s) in each journal published a median of 14 articles (IQR 8 to 25). For 1022 journals (19%), there was more than one author with the same largest number of published articles. Over the 2015–2019 period, the Gini index median was 0.183 (IQR 0.131 to 0.246), and the 95^th^ percentile was 0.355 (**Figure 2**). The correlation between the PPMP and Gini indices was 0.35 (95CI 0.33 to 0.37).

**Fig 2:**
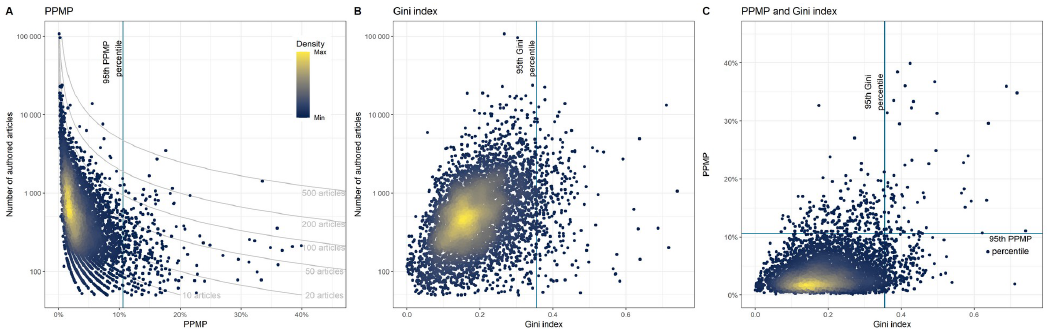
Distribution of Percentage of Papers by the Most Prolific (PPMP) author(s) (A) and Gini index (B) compared to journal size, and comparison between the PPMP and the Gini index (C), across all articles published by all journals in the United States National Library of Medicine catalog having at least one Broad Subject term and having published at least 50 authored articles between 2015 and 2019.

For both indices, there were no meaningful differences between index values across years, with the 95 ^th^ percentile ranging from 10.1% to 11.4% for the PPMP and 0.212 to 0.224 for the Gini index (**Web-Appendix 2**).

Results of the sensitivity analyses based on ‘journal articles’ alone were consistent with those for all articles (**Web-Appendix 3)**. Correlations between indices computed for all articles and for ‘journal articles’ alone were 0.94 (0.93 to 0.94) for the PPMP and 0.99 (0.99 to 0.99) for the Gini index.

In 540 journals (9.9%), for at least a quarter of the authors only the initials of their first-name(s) were presented.

#### Field-specific variations

The distribution of the PPMP and Gini index for each NLM broad term is presented in **Web-Appendix 4**. The median PPMP per field ranged from 1.1% to 9.5%, with no field having a median above the 95^th^ percentile threshold for the PPMP (i.e. 10.6%). The median Gini index per field ranged from 0.113 to 0.297, with no field having a median above the 95^th^ percentile threshold for the Gini index (i.e. 0.355).

#### Publication lag

Because of failures to report submission or acceptance dates, publication lag was not calculable for 2 743 journals (50.2%). Compared to journals that did report submission and acceptance dates, these journals had fewer authored articles (369 (IQR 200 to 712) vs 637 (IQR 355 to 1 186)). There were no differences for the Gini index but a higher PPMP (3.4% (IQR 2.0% to 5.9%) vs 2.4% (IQR 1.5% to 4.0%)). For the 2 725 journals with data on submission and publication (49.8%), the median of publication lag for all authored articles over the five years was 85 days (IQR 53 to 123) for articles published by the most prolific author(s) versus 107 days (IQR 80 to 141) for articles not published by the most prolific author(s).

**Figure 3** shows the scatter plot for all articles with the marginal density curve for the median of publication lag for the most prolific author(s) versus non-prolific authors, for each journal. For articles authored by the most prolific author(s), the distribution of the publication lag was skewed towards a shorter time-lag. Using a cut-off of 3 weeks for the median of publication lag, 277 (10.2%) of the journals had a median below this for articles by the most prolific author(s), 51 (1.9%) journals had a median below this for articles not by prolific author(s), and 38 (1.4%) journals had a median below this for both types of article (i.e authored by the most prolific author(s) or not). For the most prolific authors, publication lag decreased with the number of articles published (**Figure 4**), not solely in outlier journals. The results of the sensitivity analyses based on ‘journal articles’ alone were consistent with those for all articles (**Web-Appendix 5)**.

**Fig 3:**
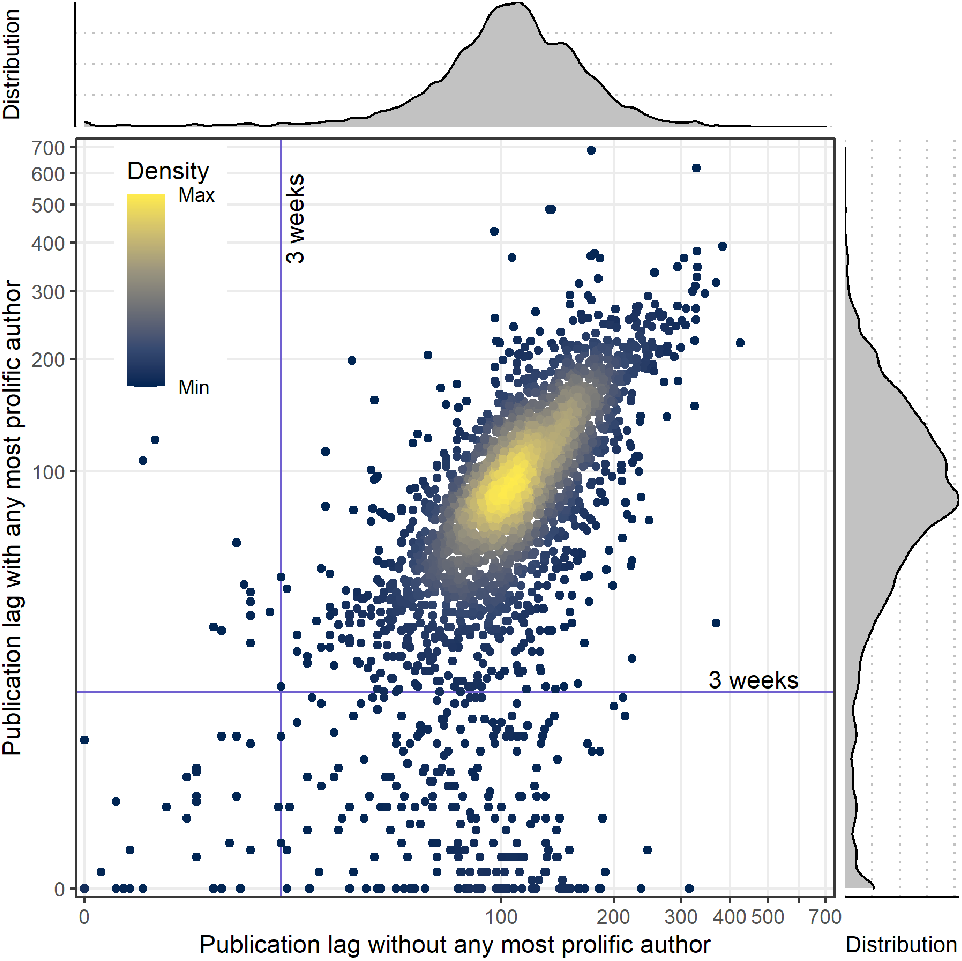
Distribution of the publication lag median across journals (in days) for articles by the most prolific authors compared to the articles without any of the most prolific authors (with marginal density plot of distributions).

**Fig 4:**
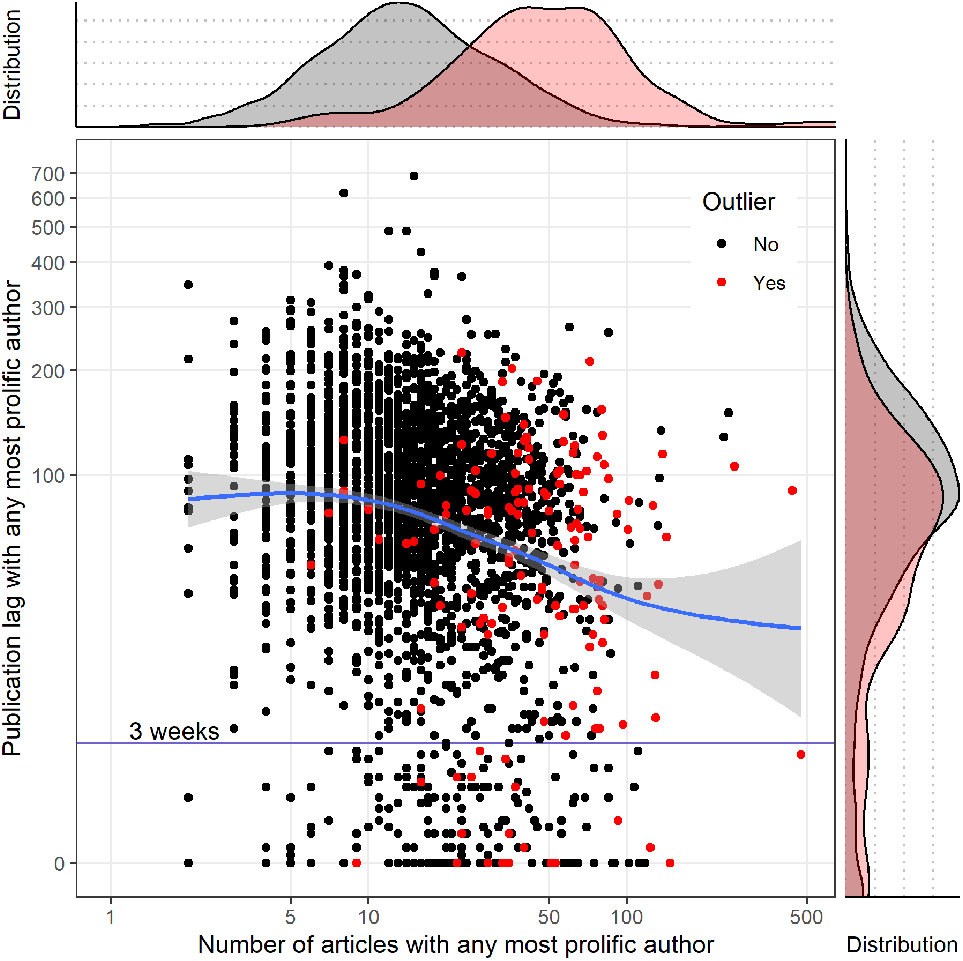
Distribution of publication lag (in days) and number of articles authored for each of the most prolific authors, across all articles (with marginal density plot of distributions).

### Description of outliers and identification of nepotistic journals

Using the 95^th^ percentile value, we identified 480 outlier journals: 206 based on the PPMP and the Gini index considered separately, and 68 based on both indices. The yearly and global distributions of these indices are presented for potential outliers as identified above (see **Web-Appendix 6**).

The main characteristics of the 100 randomly selected outlier journals are presented in **Web-Appendix 7**. Of these 100 journals, 98 were reported in English, among which 31 were also in another language (either fully multilingual journals or translation of abstracts). The most common non-English languages were German (6 journals), Chinese (5 journals), Japanese (5 journals), French and Italian (4 journals each). These outliers were well-established journals, with a median year of start of activity in 1990 (IQR 1976 to 2001).

Only 56 of these 100 journals were indexed in WoS, which enables an assessment of the journal impact factor and other citation metrics. For these 56, the median journal impact factor was 2.9 (IQR 1.5 to 4.8) with a median self-citation ratio of 0.11 (IQR 0.047 to 0.21), corresponding to a median self-citing boost according to Ioannidis and Thombs of 13% (IQR 5% to 26%).[4] The skewness and non-article inflation median was 86% (IQR 55% to 144%)[4]. Calculation of this metric was not possible for 11 journals (20%) that had a median of article citation of 0. Only 5 journals were indexed in the Directory of Open Access Journals as being full open access, and the median proportion of open access articles was 2.0% (IQR 0.47% to 8.0%). None of the outliers had an open peer review policy.

For two of the 100 journals, the full composition of the editorial boards could not be found and only the editor-in-chief was known, but was not the most prolific author. In the remaining 98 journals, at least one of the most prolific authors was a member of the editorial board in 60 journals (61%), among whom 25 (26%) were the editors-in-chief. Journals where at least one of the most prolific authors was on the editorial board tended to have a higher impact factor than the others with a median of 3.4 (IQR 2.0 to 5.3) versus a median of 1.4 (IQR 1.0 to 3.1).

Because there was sometimes more than one author with the same largest number of published articles, 108 “most prolific” authors were identified in the 100 outlier journals. We identified errors in author identification for one journal, MMW-Fortschritte der Medizin (an outlier on the Gini index), where the most prolific ‘author’ was named ‘Red’, which seems to be a diminutive for ‘Redaktion’, possibly encompassing several physical individuals. When ‘Red’ was ruled out as a valid author name, the next most prolific author was however a member of the editorial board, the PPMP increased very slightly from 7.5% of 4 920 authored articles to 7.6% of 4 553 authored articles, and the Gini index decreased very slightly from 0.637 to 0.616. Out of 1 978 identified authors (excepting ‘Red’), 1 435 (72.5%) did not publish more than one article in this journal. Among the remaining 107 individual authors, 95 were formally identified from Web of Science. Among the 12 remaining, identification was considered unreliable for 8 because of possible homonyms and 4 were not indexed in WoS. For these 12 authors, manual disambiguation using PubMed and Google identified 6 journalists, 5 physicians with a consistent affiliation to the journal considered, and one author where a high risk of homonym persisted (‘Wang, Y’ in *Journal of clinical otorhinolaryngology, head, and neck surgery*). Among the 95 other authors, the median of the H-index was 28 (IQR 13 to 50).

## Discussion

In this comprehensive survey of 5 468 biomedical journals, we characterized several features of editor-author relationships among which were the following: (i) article output was sometimes dominated by the prolific contribution of one author or a group of authors, (ii) time lags to publication were in some instances shorter for these prolific authors and (iii) prolific authors were typically members of the journal’s editorial board.

We concluded that defining the top 5% nepotistic journals required the threshold to be set at up to 10.6% of articles published by the most prolific author. In absolute terms, we believe it is reasonable to question the judgement of an editor where more than 10% of the published papers are authored by the same person. To better characterize the lack of heterogeneity in authorship, we also computed the Gini index, for which the corresponding 95th percentile was 0.355 over five years and 0.20 when computed annually. This suggests that a broader time-frame allowed for more occasional authors to be recruited while maintaining the regular authors, revealing the latent heterogeneity.

The PPMP and the Gini index explore complementary patterns of asymmetry in publishing patterns. The PPMP reflects author practices while the Gini is more sensitive to groups of highly prolific authors. The Gini index has one advantage over the PPMP, in that it is less constrained by the total number of articles published. If a journal publishes a very large number of papers, it becomes increasingly implausible that a single prolific author could account for 10% or more of them, as there is a natural upper limit to how many papers any one individual can author. Conversely, for a journal that publishes very few papers, an author could be identified as above the 95th centile with a relatively modest number of publications.

For the subgroup of journals reporting submission and publication dates, the time-lag for the most prolific author(s) was shorter, suggesting that for certain journals, that are outliers on PPMP and/or Gini, peer reviews may have been absent or only superficial for prolific authors. However, for the most prolific authors, the publication lag decreases with the number of articles published across all journals and not solely in the sub-sample of outliers. This suggests that our description of these outliers based on PPMP could be only the tip of the iceberg, capturing solely the most extreme cases of hyper-prolific publication in a given journal.

Our findings persisted when all articles were considered as well as when only ‘journal articles’ were considered (i.e. excluding articles explicitly referenced as editorial, correspondence or news articles), suggesting that editorials, correspondence and news, are not the only drivers of the indicators we explored. Conversely, it is possible that the labelling as ‘journal article’ is not perfect and carries a risk of misclassification bias.

We should beware of assuming that a hyper-prolific author is necessarily engaged in questionable publishing practices: some people are naturally highly productive, and the speed with which good research can be completed is highly variable across research fields. Furthermore, authors may be represented in many papers because they play a key role in one aspect of the research, such as the statistical analysis. Similarly, shorter publication lags may occur simply because it is easier to find reviewers for eminent authors, or in a particular subject area, and/or because their expertise means that their papers require less revision. Nevertheless, there is no doubt that some highly prolific authors achieve an unusual level of productivity by exploiting the system or engaging in academic misconduct.[5,6] It is important to make a distinction between hyper-prolific authors who publish a lot in a range of different journals and those who are exploiting a select pool of a few journals in which they appear as prominent authors (as we explored here). It is also very important to complement the PPMP and the Gini index with the absolute numbers of papers authored by the most prolific authors, because some problematic journal behaviours could pass unnoticed when only these two indices are used.

On a random sample of outlier journals identified using the PPMP and/or the Gini index, we found that the prolific authors can be ‘established’ scientists with a relatively high h-Index (e.g. a median of 28), and that 60% of these most prolific authors were editors-in-chief or members of the journal’s editorial board. About half of these journals had a median journal impact factor of 2.9 (IQR 1.5 to 4.8). These journals generally presented a large self-citation ratio, meaning that some of them may have questionable practices by manipulating their impact factor. The other half of these journals did not have an impact factor, possibly indicating they were new journals joining the WoS. Even though WoS uses an extensive list of eligibility criteria, it is also possible that some of the new journals are predatory, journals that are known to have “leaked into” trusted sources. [7]^7^

Our results underscore possible problematic relationships between authors who sit on editorial boards and decision-making editors. Typically, publishers promote independence between authors and journals. Hyper-published authors may see such relationships as a way to more easily reach publication thresholds for hiring, promotion and tenure. There may be defensible reasons for members of the editorial board of a journal to hyper-publish in a journal.[2] There are for instance certain research fields that are research niches, where the contributing authors are part of a very small community of specialists and are therefore the most likely authors.

Although our findings are based solely on a sub-sample of journals, they provide crucial evidence that editorial decisions were not only unusually, but also selectively, fast for the favoured subset of prolific authors. This pattern was also found by Sarigöl et al. when exploring favouritism towards collaborators and co-editors, which persists even after taking into account individual article quality, measured as citation and download numbers.[8] This phenomenon could have an impact on productivity-based metrics and suggests a risk of instrumentalisation, if not corruption, of the scientific enterprise, by using journals as a ‘publication laundry’ for ‘vanity publication’[9] by authors closely related to the editorial board. Exploiting productivity metrics has been widely described, in the form of self-citation, honorary authorship and/or ghost writing. Manipulation of individual metrics by resorting to a dedicated ‘nepotistic’ journal appears to be a little studied way of exploiting the system.

### Limitations

Our descriptive and exploratory survey, based on a large available database, provides information about the broad scene of ‘nepotistic journals’ but it may miss some finer points, especially concerning the quality of articles published in these journals. The quality of a scientific article is a difficult concept to measure and it cannot be easily summarized in quantitative metrics. In addition, we restricted our analysis to journals indexed in the National Library of Medicine (NLM) under one or more of the existing broad terms. Some journals are registered without broad terms, requiring a manual pick-up by the NLM. Consequently, our survey may have preferentially included the more established journals indexed in the NLM, with a durable presence in the database, and hence likely to have a better-quality global and editorial conduct than non-indexed journals. Similarly, because we restricted our search to journals publishing a minimum of 50 papers in the 2015-2019 period, we may have missed smaller journals with less professional editorial practices.

Importantly, our automated calculations carry a risk of inaccuracy as a result of homonymy. Misidentification and/or merging of author names could bias the PPMP and the Gini index in both directions, and the risk of merging increases when only the initial or first-name is known, and in case of authors with similar names. The greater risk of homonyms could partially explain the increased Gini index values for larger journals, without reference to a tendency to editorial misconduct. Our analysis of the random sample of outliers enabled a disambiguation procedure consisting in inspecting qualitatively the most prolific authors. Only 1 out of 108 “most prolific” authors within a given journal was considered to be at being at high risk of homonymy. Among these 108 authors, this procedure also enabled identification of the 6 most prolific authors, who were professional journalists for whom high productivity is of course not an indicator of any academic misconduct, as they are professionals paid by the journal and not academics. The two proposed indicators, and their current calculation, should therefore not be used indiscriminately but could rather serve as a screening tool for potentially problematic journals that may then require careful exploration of their editorial practices.

While our results are exploratory and do not yet support a widespread use of these indicators, we hope that further research will help to establish these easily computed indexes as a resource for publishers, authors, and indeed scientific committees involved in promotion and tenure, to screen for potentially biased journals needing further investigation. DORA paved the way, of moving away from productivity-based metrics, and other efforts followed such as the Hong Kong Principles for assessing researchers. Integrity-based metrics are indeed needed to overcome the limitations of productivity-based metrics [10]. A transparent declaration of interests in communicating research is surely one important aspect of scientific integrity and trustworthy science. This principle of course applies to financial conflicts of interest which are often under-declared by journal editors,[11] and also to non-financial conflicts of interest such as editor-author relationships.

The proposed indices could add transparency in the editorial decision-making and peer review process of any journal. This transparency is currently lacking towards the public and any stakeholder involved in the research community, such as COPE, the Committee for Publication Ethics. Guidance for editors and publishers should be developed to delineate good practices and prevent obvious misconduct.

## Methods

We developed and followed a research protocol, which was prospectively registered on 21st July 2020 on the Open Science Framework (https://osf.io/6e3uf/). The analytic code and summarized data are also available on the same URL.

### Data extraction

The eligibility criteria for the selection of journals were (i) a biomedical journal referenced in the National Library of Medicine (NLM), (ii) having at least one ‘Broad Subject Term’ and, (iii) having published more than 50 papers between January 2015 and December 2019. Broad Subject Terms are Medical Subject Headings (MeSH), terms used to describe a journal’s overall scope, and they are defined by the NLM for journals in the MEDLINE database.[12] The 2015–2019 period was chosen, as this 5-year window enables a smoothing of random variations and description of recent practices. One author (AS) searched for changes to journal names during the 2015–2019 period and, in cases of renaming, pooled the articles published under the different names.

To identify eligible journals, we used the Entrez programming utilities (E-utilities) which enable queries to the National Center for Biotechnology Information (NCBI) databases. The search query – presented in **Web-Appendix 8** – was used to identify all biomedical journals in the NLM catalogue having at least one of the ‘Broad Subject Terms’ listed. Then, for each journal, article metadata was automatically collected with E-utilities. On account of technical restrictions, querying for article metadata was run from 2015 up to the date of extraction and then restricted to the period January 2015 to December 2019.

To manage articles without an author name, the third selection criterion was slightly modified to focus on journals with at least 50 ‘authored articles’ – i.e. articles with at least one identified author – over the 2015–2019 period (see ‘protocol changes’). Publications reprinted in several journals (e.g. PRISMA statements published in 6 different journals to promote dissemination) did not receive special treatment, and no correction was applied, as each article was only examined in relation to its publication journal.

### Index calculation

#### Percentage of papers by the most prolific author and Gini index

For each journal, each author was identified by his or her full name (i.e. family name and complete first name) or barring that, by his or her family name and first name initial(s). The number of articles authored by this person was counted. When there was more than one author with the same largest number of published articles, they were all considered as the “most prolific” authors. The PPMP was defined as the number of articles by the most prolific author (n_max_) divided by the total number of authored articles in the journal (N_tot_): PPMP = [n_max_ / N_tot_]. Complementary to the PPMP, the Gini index was used to explore inequality in the number of published articles related to more than one author. The Gini index for the number of publications by each author was calculated, with correction for the total number of authors (see formula and example in **Web-Appendix 9**)[3]. The values of the Gini index range from 0 (perfect equality in numbers of articles among authors) to 1 (major inequality).

For the primary analysis, these two outcomes (PPMP and Gini) were computed for all papers, and for a sensitivity analysis they were computed only for papers labelled as ‘journal articles’ (using the NCBI publication type).

#### Publication time-lag

For each article, the publication lag – defined as the time between submission and acceptance of an article – was computed whenever possible. After this, each journal was characterized by (i) median publication lag for articles authored by at least one of the most prolific authors and (ii) median publication lag for articles not authored by the most prolific author(s).

### Description of outlier journals

Outliers were defined as journals with a PPMP value and/or the Gini index above their respective 95 ^th^ percentiles in the principal analysis (i.e. on all articles). For pragmatic reasons a sample of 100 outlier journals was randomly selected (first sorted by full name, in alphabetic order, and randomly sampled using a random number generator with a seed arbitrarily set at 42; R function *sample_n* in *dplyr* package). One reviewer (AS or FN) manually extracted characteristics related to the journal impact factor (Web of Science: WoS), open access policies (WoS and Directory of Open Access Journals), open peer review policies (Publons - Clarivate), the most prolific authors’ h-index (WoS) and presence and role (i.e. editor in chief or board member) on the editorial board of the journal (journal or publisher website). Where this information was available, we made a distinction between advisory boards (who were not considered) and editorial boards. For the year 2019, the metrics ‘self-citation boost’ (i.e. number of self-citing articles over number of non-self-citing articles) and ‘skewness & non-article inflation’ (i.e. impact factor minus median of citations for an article, over median of citations for an article) was computed according to Ioannidis and Thombs[4].

### Data analysis

A descriptive analysis was performed using median, range, and quartiles for continuous variables, and counts and percentages for categorical variables. Descriptions for the 100 outlier journals were computed overall and with respect to membership of any of the most prolific author(s) on the editorial board. Correlations were computed using Pearson’s coefficient, with 95% confidence interval (CI).

To explore field-specific variations, the distribution of the two indices within each ‘Broad Subject term’ was graphically displayed. The yearly and overall distribution of the percentage of papers by each author and the Lorenz curve – a graphic representation of the cumulative distribution of appearances as an author – were presented for each of the potential outliers identified above.

All analyses were conducted using R version 3.6,[13] and main packages RISmed 2.1 for queries on journal characteristics,[14] easyPubMed 2.13 for queries on article characteristics,[15] DescTools 0.99 for Gini index calculation,[16] and tidyverse 1.3 for miscellaneous.[17]

### Protocol changes

Some practical challenges arose in our research because some articles unexpectedly lacked author names, which precluded them from contributing to the numerator of PPMP or to the Gini index. We therefore amended our definitions to make it explicit that the PPMP denominator was defined as the number of articles with at least one identified author rather than all published articles, and journals were included only if they had published 50 articles with author names rather than all published articles.

The 3-week threshold used to describe publication lag as being suggestive of unduly rapid or absent peer review was not initially specified in our protocol, and was added for descriptive purposes. We also explored the relationship between publication lag for articles authored by any of the most prolific author (s) and the number of papers authored by these authors. The description of the outlier journals with respect to the membership of any of the most prolific author(s) on the editorial board was added *a posteriori* for exploratory purposes.

## Contributions

C.L., F.N. and A.S. initiated the project and drafted the protocol. All authors contributed and approved the protocol. F.N. and A.S. extracted the data. A.S. wrangled with the data. C.L., F.N. and A.S. wrote the first drafted the completed study to which all authors revised and approved final submitted version of the article.

## Web-Appendixes Web-Appendix 1

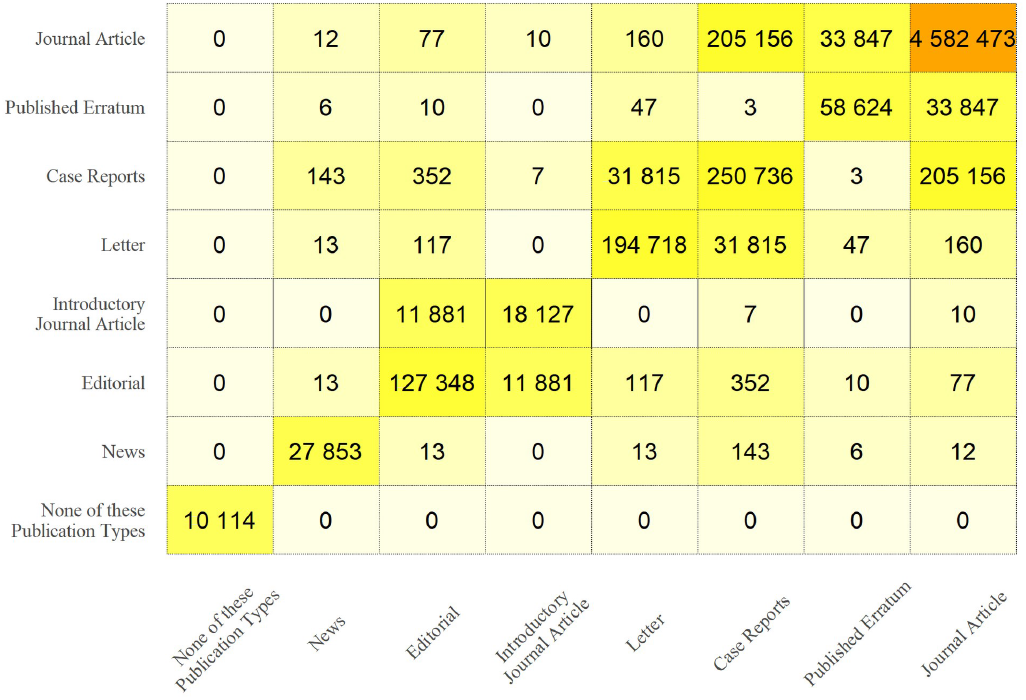

Publication types attached to each article and their co-occurences, among all journals in the United States National Library of Medicine catalog having at least one Broad Subject term and having published at least 50 signed articles between 2015 and 2019.

## Web-Appendix 2

|  |  | All articles |  |  |  |  | Journal Article |  |  |  |  |
| --- | --- | --- | --- | --- | --- | --- | --- | --- | --- | --- | --- |
| Year |  | 2015 | 2016 | 2017 | 2018 | 2019 | 2015 | 2016 | 2017 | 2018 | 2019 |
| Number of journals with ≥ 50 authored articles |  | 4149 | 4143 | 4100 | 4065 | 4131 | 3994 | 4031 | 3977 | 3961 | 4020 |
| Number of articles | Median |  |  |  |  |  |  |  |  |  |  |
|  | [IQR] | 130 [82 - 231] | 131 [83 - 230] | 132 [85 - 235] | 136 [85 - 242] | 142 [87 - 251] | 123 [80 - 214] | 123 [80 - 213] | 124 [81 - 221] | 128 [81 - 226] | 133 [83 - 232] |
|  | Range | 50 - 29 762 | 50 - 23 021 | 50 - 25 330 | 50 - 18 832 | 50 - 20 423 | 50 - 28 108 | 50 - 22 142 | 50 - 25 322 | 50 - 18 832 | 50 - 19 877 |
| Number of articles with an author |  |  |  |  |  |  |  |  |  |  |  |
|  | Median |  |  |  |  |  |  |  |  |  |  |
|  | [IQR] | 128 [82 - 228] | 129 [82 - 227] | 130 [83 - 233] | 134 [84 - 240] | 140 [86 - 249] | 122 [79 - 213] | 121 [79 - 211] | 122 [80 - 218] | 127 [80 - 223] | 132 [82 - 230] |
|  | Range | 50 - 28 937 | 50 - 22 827 | 50 - 25 327 | 50 - 18 635 | 50 - 20 423 | 50 - 28 094 | 50 - 22 120 | 50 - 25 319 | 50 - 18 635 | 50 - 19 877 |
| PPMP (%) | Median |  |  |  |  |  |  |  |  |  |  |
|  | [IQR] | 3.75 [2.37 - 5.80] | 3.67 [2.41 - 5.77] | 3.66 [2.37 - 5.71] | 3.63 [2.33 - 5.56] | 3.57 [2.27 - 5.47] | 3.53 [2.29 - 5.46] | 3.54 [2.34 - 5.45] | 3.53 [2.31 - 5.29] | 3.49 [2.25 - 5.26] | 3.42 [2.20 - 5.26] |
|  | Range | 0.2 - 39.2 | 0.1 - 46.3 | 0.1 - 50.4 | 0.1 - 52.1 | 0.1 - 44.4 | 0.2 - 41.2 | 0.1 - 43.9 | 0.2 - 59.2 | 0.1 - 55.0 | 0.1 - 50.6 |
| PPMP 95th percentile (%) |  | 11.4 | 10.8 | 10.4 | 10.1 | 10.4 | 10.0 | 10.2 | 9.84 | 9.62 | 9.80 |
| Number of articles by MPA |  |  |  |  |  |  |  |  |  |  |  |
|  | Median |  |  |  |  |  |  |  |  |  |  |
|  | [IQR] | 5 [4 - 8] | 5 [3 - 8] | 5 [3 - 8] | 5 [4 - 8] | 5 [4 - 8] | 5 [3 - 7] | 5 [3 - 7] | 5 [3 - 7] | 5 [3 - 7] | 5 [3 - 7] |
|  | Range | 1 - 195 | 1 - 172 | 1 - 194 | 1 - 161 | 1 - 226 | 1 - 109 | 1 - 88 | 1 - 193 | 1 - 140 | 1 - 217 |
| Tied as MPA |  | 1635 (39%) | 1623 (39%) | 1631 (40%) | 1655 (41%) | 1616 (39%) | 1774 (44%) | 1733 (43%) | 1804 (45%) | 1811 (46%) | 1754 (44%) |
| Gini index | Median | 0.087 [0.055 - | 0.086 [0.054 - | 0.085 [0.054 - | 0.085 [0.055 - | 0.085 [0.055 - | 0.077 [0.047 - | 0.077 [0.048 - | 0.078 [0.048 - | 0.077 [0.050 - | 0.076 [0.047 - |
|  | [IQR] | 0.132] | 0.128] | 0.127] | 0.127] | 0.129] | 0.119] | 0.117] | 0.117] | 0.116] | 0.115] |
|  | Range | 0.00 - 0.67 | 0.00 - 0.68 | 0.00 - 0.66 | 0.00 - 0.76 | 0.00 - 0.63 | 0.00 - 0.67 | 0.00 - 0.60 | 0.00 - 0.64 | 0.00 - 0.76 | 0.00 - 0.64 |
| Gini index 95th percentile |  | 0.222 | 0.224 | 0.215 | 0.212 | 0.216 | 0.210 | 0.213 | 0.202 | 0.201 | 0.203 |
| Median of publication lag ratio (MPA / no MPA) |  |  |  |  |  |  |  |  |  |  |  |
|  | Median |  |  |  |  |  |  |  |  |  |  |
|  | [IQR] | 0.84 [0.55 - 1.12] | 0.84 [0.57 - 1.12] | 0.85 [0.56 - 1.10] | 0.83 [0.55 - 1.13] | 0.86 [0.59 - 1.14] | 0.85 [0.59 - 1.12] | 0.87 [0.61 - 1.14] | 0.86 [0.60 - 1.11] | 0.86 [0.61 - 1.14] | 0.89 [0.64 - 1.14] |
|  | Range | 0.0 - 35.0 | 0.0 - 79.0 | 0.0 - 28.5 | 0.0 - 24.8 | 0.0 - 92.5 | 0.0 - 35.0 | 0.0 - 79.0 | 0.0 - 4.0 | 0.0 - 16.7 | 0.0 - 7.7 |
| Not calculable |  | 2420 (58%) | 2295 (55%) | 2204 (54%) | 2178 (54%) | 2243 (54%) | 2335 (58%) | 2271 (56%) | 2163 (54%) | 2163 (55%) | 2229 (55%) |

## Web-Appendix 3

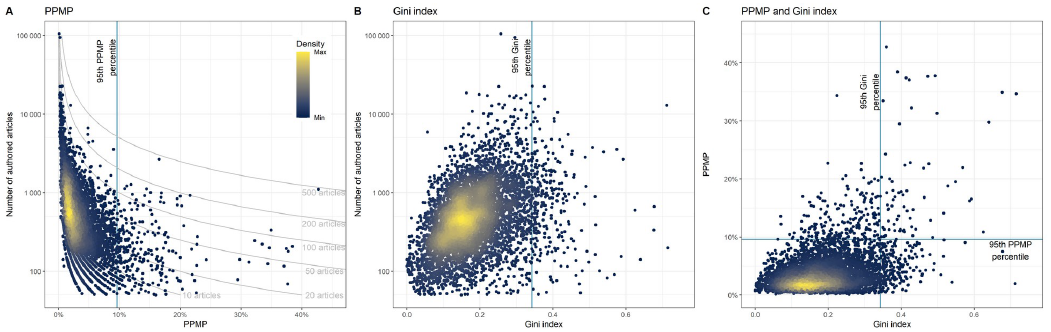

Distribution of the Percentage of Papers by the Most Prolific (PPMP) author(s) (A) and the Gini index (B) in relation to journal size, and comparison between the PPMP and the Gini index (C), among articles labelled as ‘Journal Articles’ only, published by all journals in the United States National Library of Medicine catalog having at least one Broad Subject term and having published at least 50 signed articles between 2015 and 2019.

## Web-Appendix 4

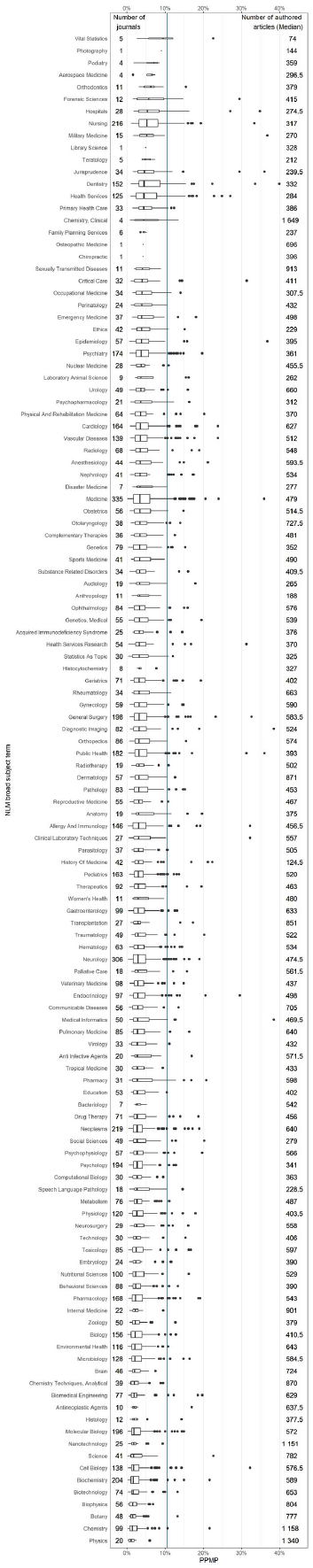

A) Distribution of the Percentage of Papers by the Most Prolific (PPMP) author for each United States National Library of Medicine (NLM) broad term, for NLM represented by at least 10 journals having published at least 50 signed articles between 2015 and 2019. The number of journal covered by a Broad Subject term is shown next to the name of the field of study. Width of the box-and-whisker relative to the number of journals. Vertical line at the 95^th^ percentile of PPMP among journals.

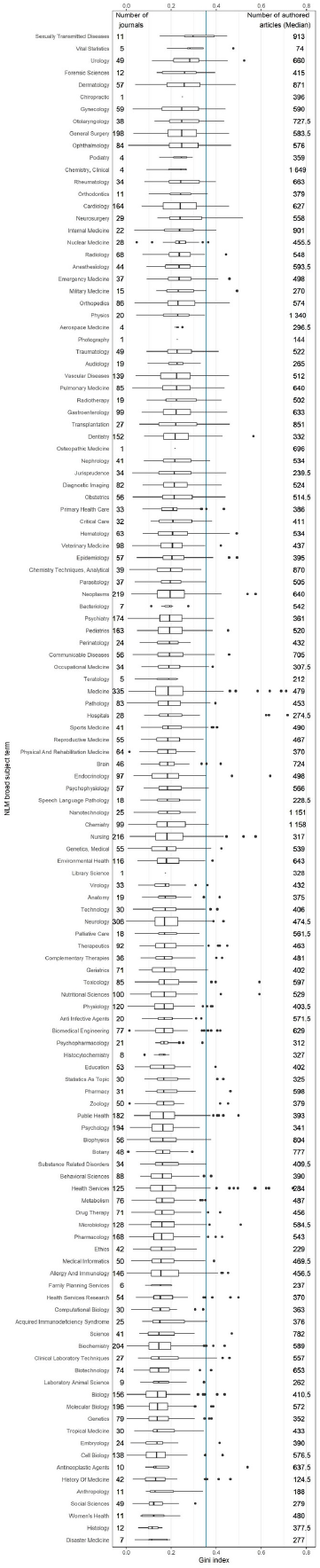

B) Distribution of the Gini index for each United States National Library of Medicine (NLM) broad term, for NLM represented by at least 10 journals having published at least 50 signed articles between 2015 and 2019. The number of journal covered by a Broad Subject term is shown next to the name of the field of study. Width of the box-and-whisker relative to the number of journals. Vertical line at the 95^th^ percentile of PPMP among journals.

## Web-Appendix 5

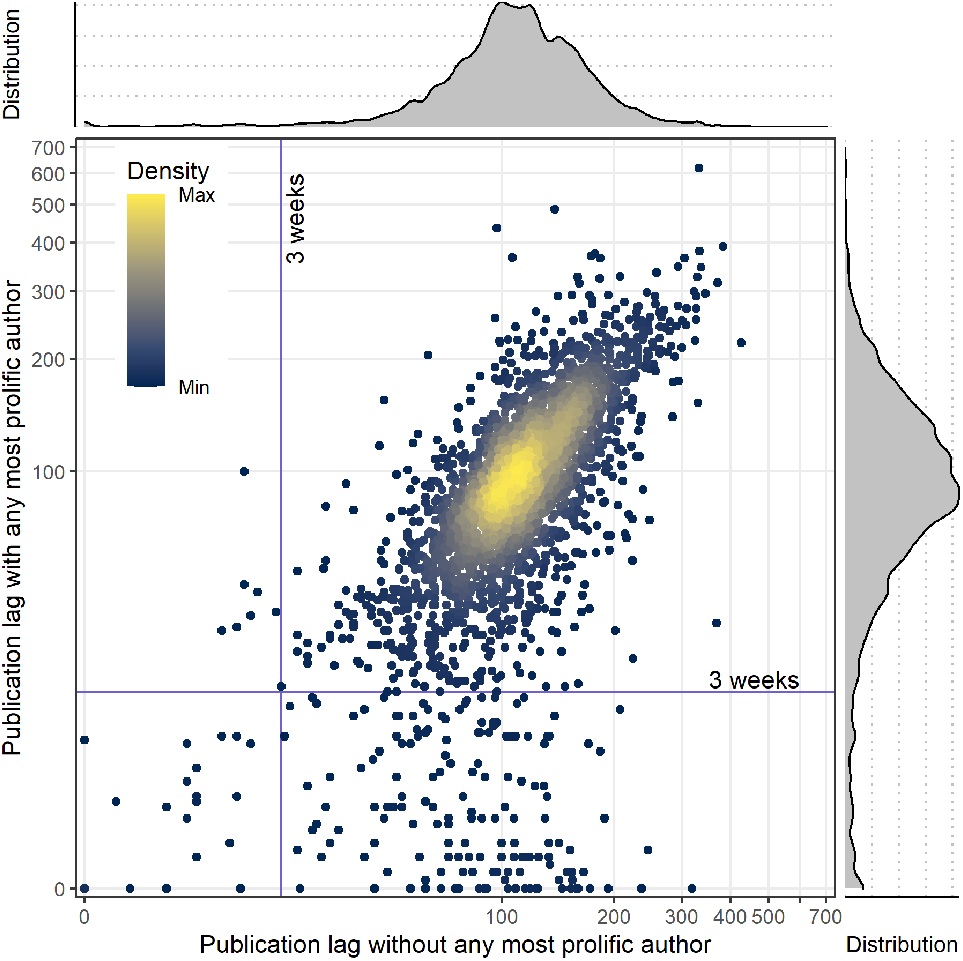

A) Distribution of the publication lag median across journals (in days) for articles by the most prolific authors compared to the articles without any of the most prolific authors, among articles labelled as ‘‘Journal Articles” only (with marginal density plot of distributions).

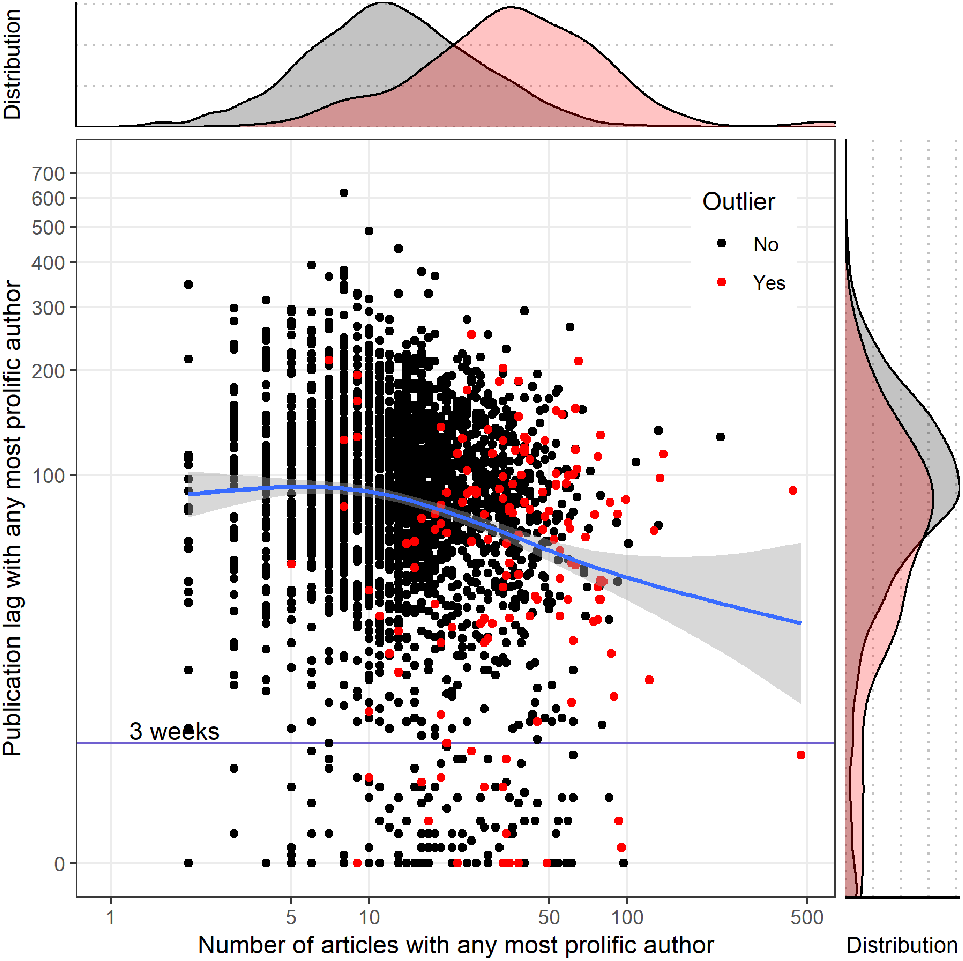

B) Distribution of publication lag (in days) and number of articles signed for each of the most prolific authors, across all articles, among articles labelled as ”Journal Articles” only (with marginal density plot of distributions).

## Web-Appendix 6

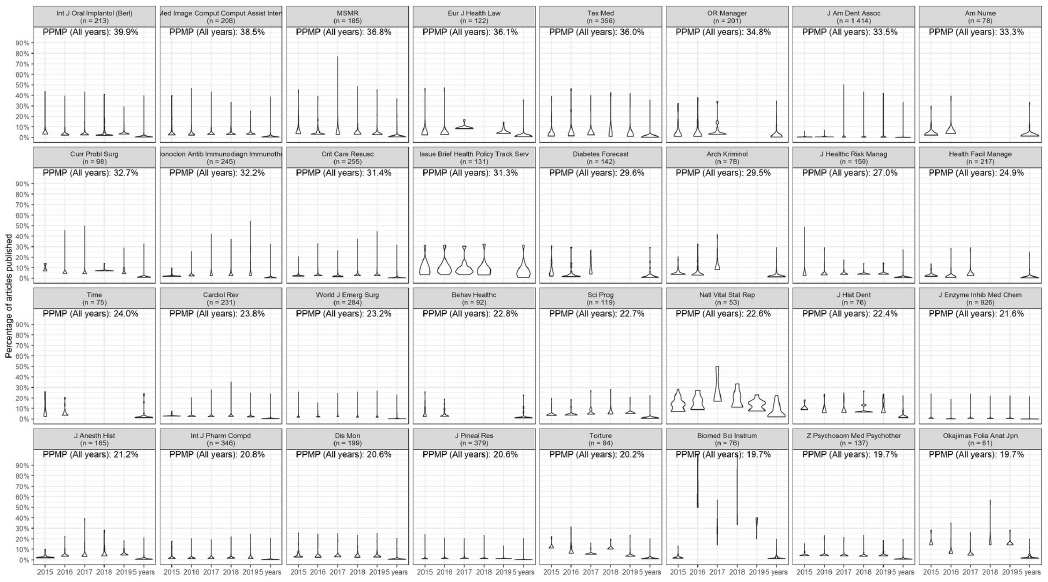

A) Yearly and global distribution of the percentage of papers by each author for each outlier journal identified as having a Percentage of Papers by the Most Prolific (PPMP) author or a Gini index reaching the 95^th^ percentile or over for journals having at least one NLM broad subject term, for the period 2016 to 2019.

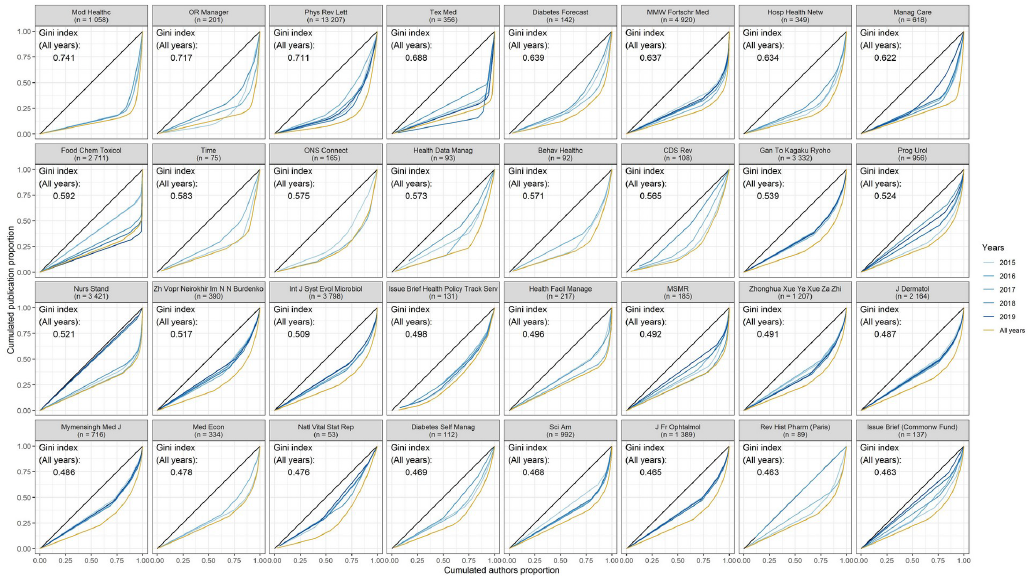

B) Yearly and global Lorenz distribution curve for number of publications by each author, for each outlier journal identified as having a Percentage of Papers by the Most Prolific (PPMP) author or a Gini index reaching the 95^th^ percentile or over for journals having at least one NLM broad subject term for the period 2016-2019. The Lorenz curve is a visual representation of non-homogeneity in the distribution of authorship, with a straight line between 0 and 1 representing a perfect equality. The Gini index is the area between the line of equality and the Lorenz curve.

## Web-Appendix 7

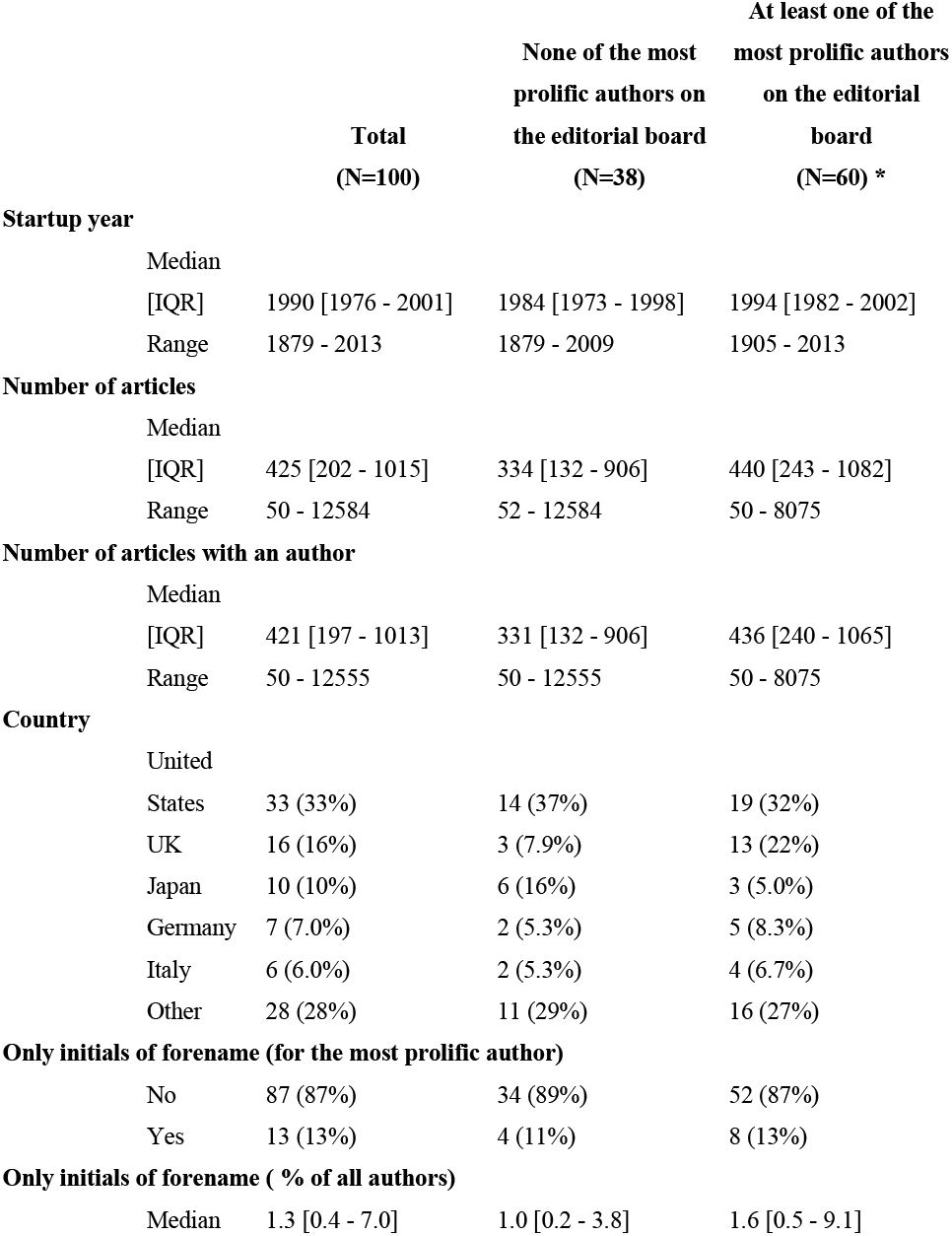
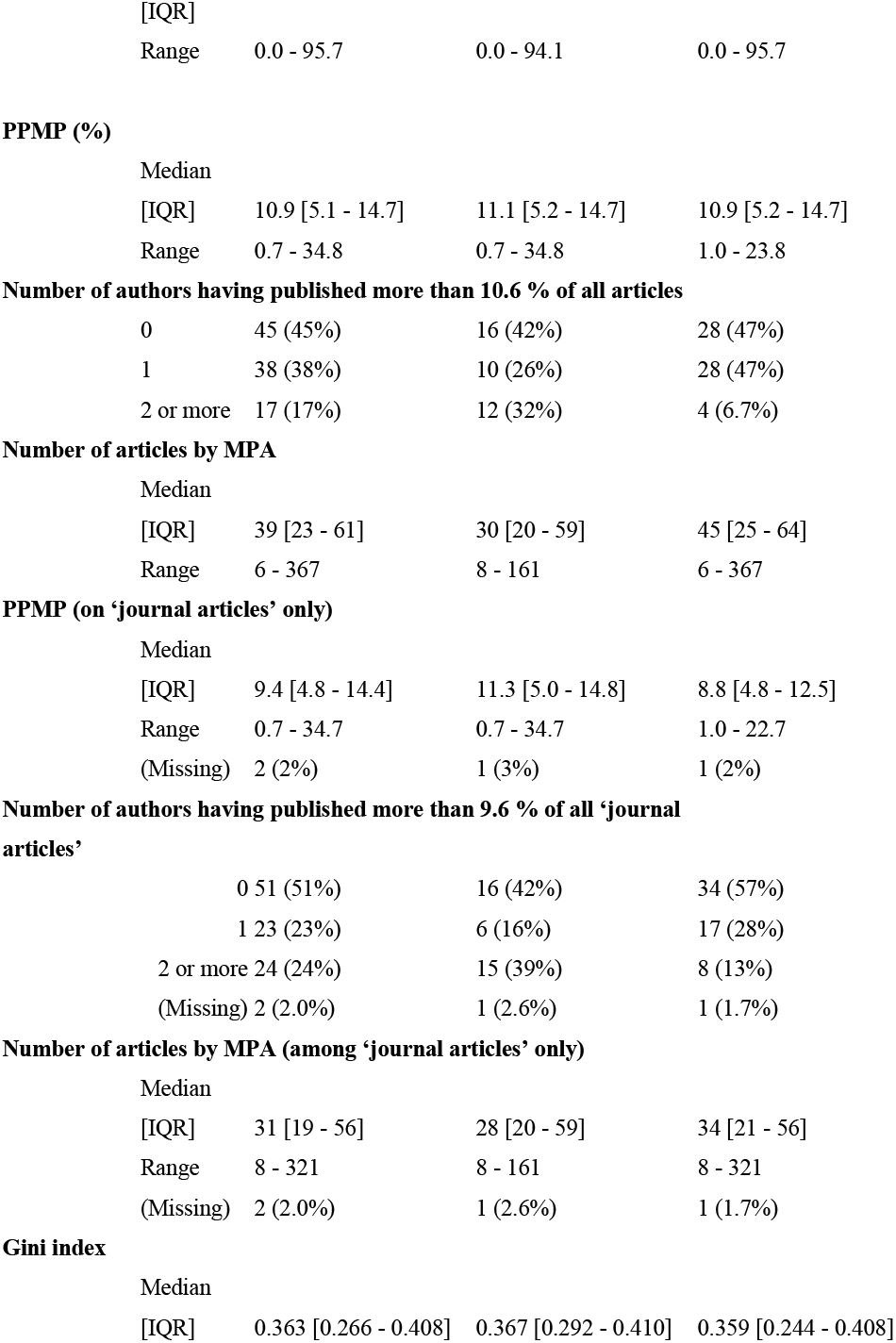
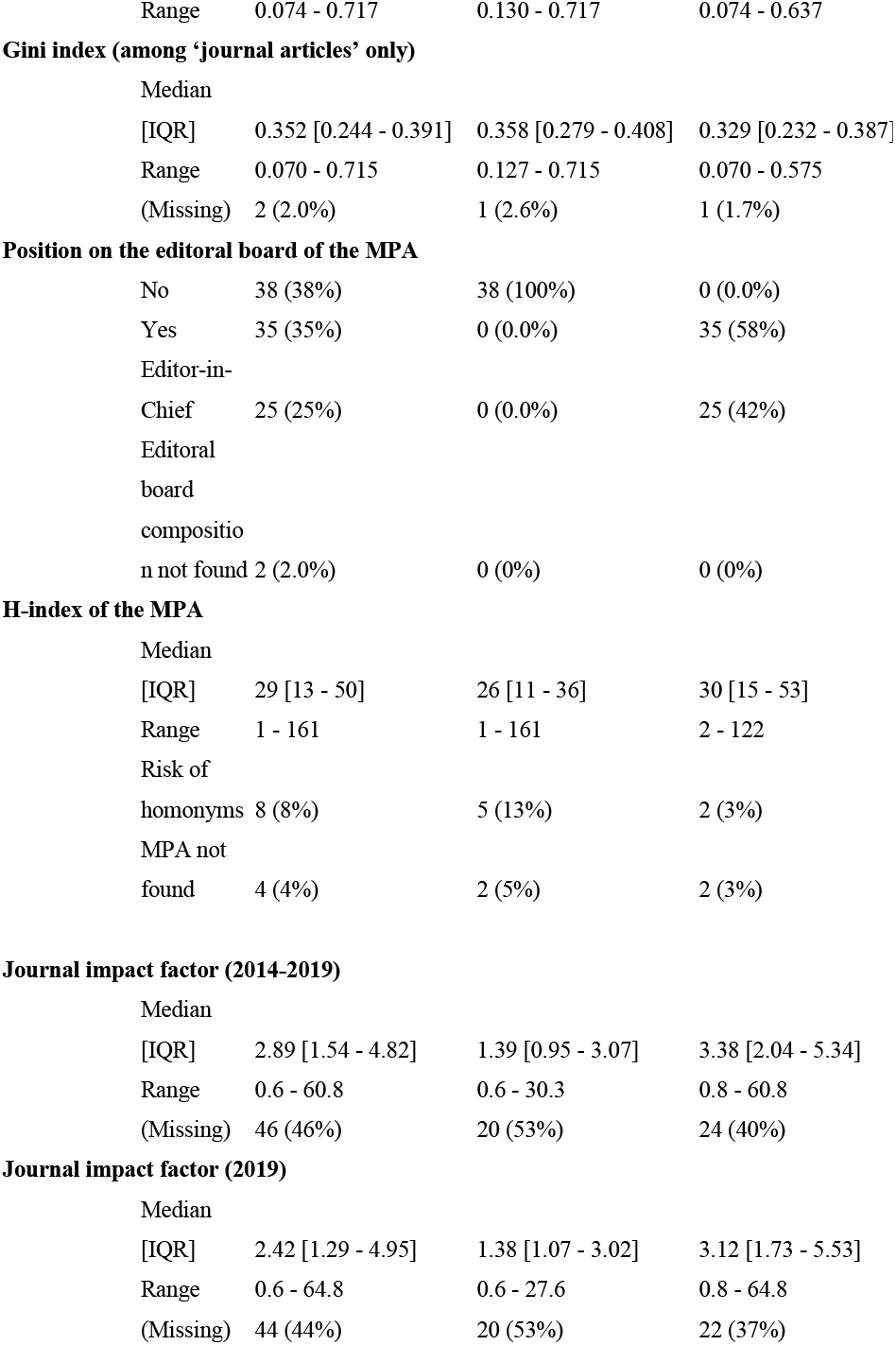
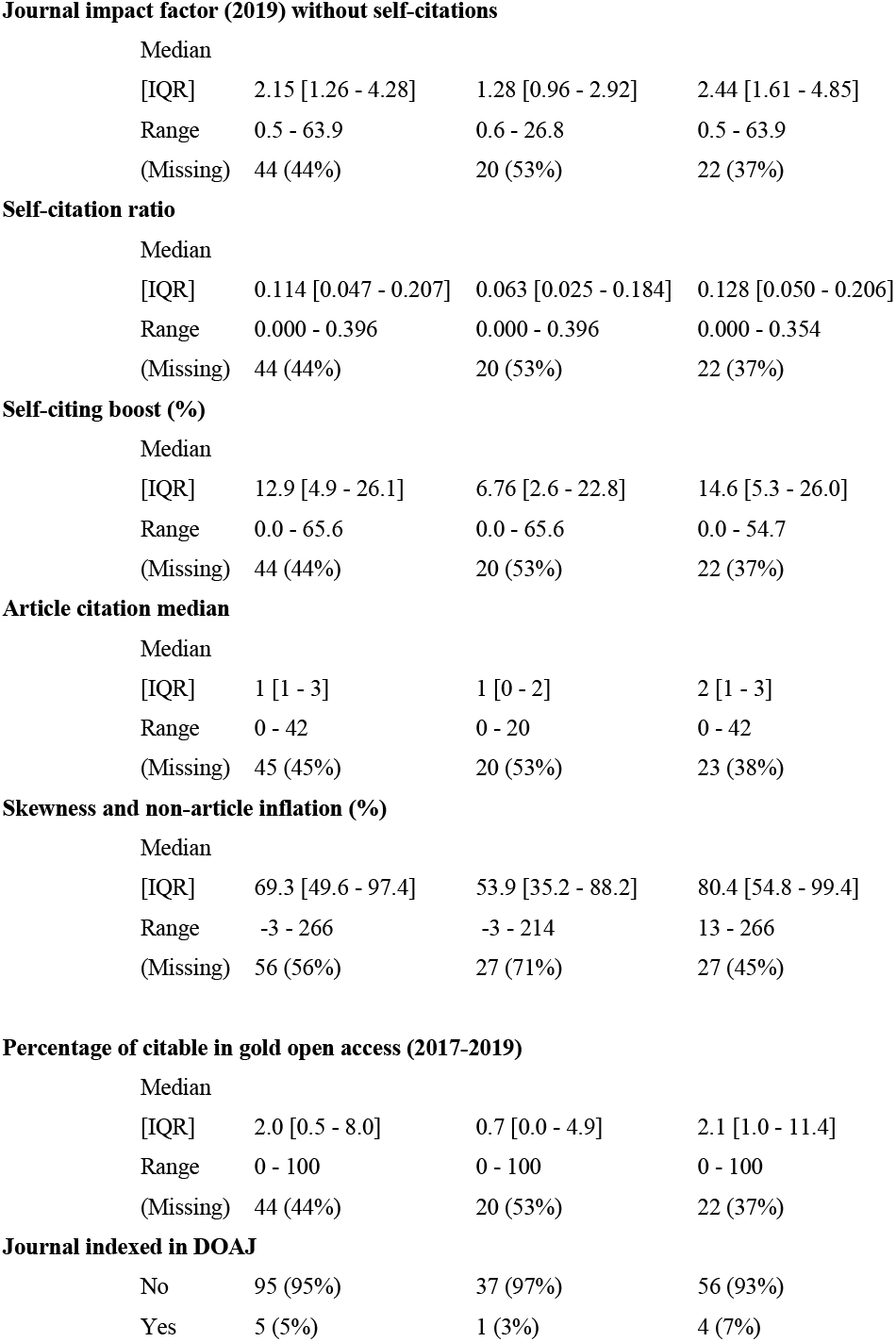
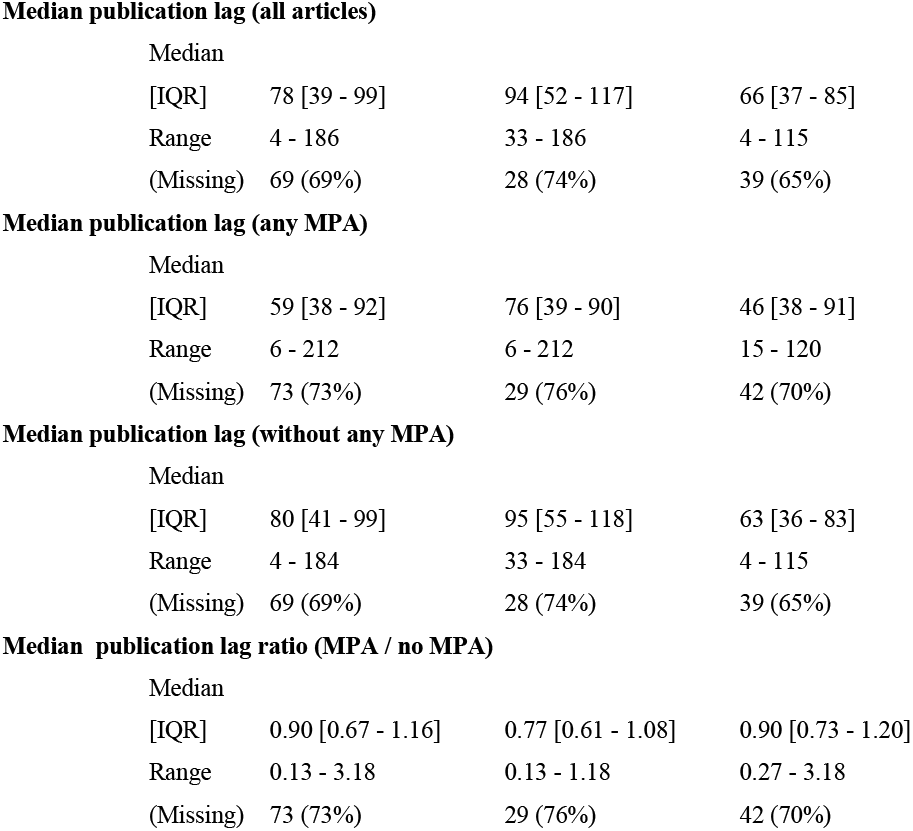

Main characteristics of 100 randomly selected journals with a PPMP or a Gini index higher than the 95 ^th^ percentile, among the journals in the United States National Library of Medicine catalog having at least one Broad Subject term and having published at least 50 signed articles between 2015 and 2019. * For two journals, the position on the editorial board of the most prolific author could not be established.

## Web-Appendix 8

United States National Library of Medicine catalog journal identification query:

(“acquired immunodeficiency syndrome”[st] OR “aerospace medicine”[st] OR “allergy and immunology”[st] OR “anatomy”[st] OR “anesthesiology”[st] OR “anthropology”[st] OR “anti infective agents”[st] OR “antineoplastic agents”[st] OR “audiology”[st] OR “bacteriology”[st] OR “behavioral sciences”[st] OR “biochemistry”[st] OR “biology”[st] OR “biomedical engineering” [st] OR “biophysics”[st] OR “biotechnology”[st] OR “botany”[st] OR “brain”[st] OR “cardiology”[st] OR “cell biology”[st] OR “chemistry”[st] OR “chemistry techniques, analytical”[st] OR “chemistry, clinical”[st] OR “chiropractic”[st] OR “clinical laboratory techniques”[st] OR “communicable diseases”[st] OR “complementary therapies”[st] OR “computational biology”[st] OR “critical care”[st] OR “dentistry”[st] OR “dermatology”[st] OR “diagnostic imaging”[st] OR “disaster medicine”[st] OR “drug therapy”[st] OR “education”[st] OR “embryology”[st] OR “emergency medicine”[st] OR “endocrinology” [st] OR “environmental health”[st] OR “epidemiology”[st] OR “ethics”[st] OR “family planning services”[st] OR “forensic sciences”[st] OR “gastroenterology”[st] OR “general surgery”[st] OR “genetics”[st] OR “genetics, medical”[st] OR “geriatrics”[st] OR “gynecology”[st] OR “health services”[st] OR “health services research”[st] OR “hematology”[st] OR “histocytochemistry”[st] OR “histology”[st] OR “history of medicine”[st] OR “hospitals”[st] OR “internal medicine”[st] OR “jurisprudence”[st] OR “laboratory animal science”[st] OR “library science”[st] OR “medical informatics”[st] OR “medicine”[st] OR “metabolism”[st] OR “microbiology”[st] OR “military medicine”[st] OR “molecular biology”[st] OR “nanotechnology”[st] OR “neoplasms”[st] OR “nephrology”[st] OR “neurology”[st] OR “neurosurgery”[st] OR “nuclear medicine”[st] OR “nursing”[st] OR “nutritional sciences”[st] OR “obstetrics”[st] OR “occupational medicine”[st] OR “ophthalmology”[st] OR “orthodontics”[st] OR “orthopedics”[st] OR “osteopathic medicine”[st] OR “otolaryngology”[st] OR “palliative care”[st] OR “parasitology”[st] OR “pathology”[st] OR “pediatrics”[st] OR “perinatology”[st] OR “pharmacology”[st] OR “pharmacy”[st] OR “photography”[st] OR “physical and rehabilitation medicine”[st] OR “physics”[st] OR “physiology”[st] OR “podiatry”[st] OR “primary health care”[st] OR “psychiatry”[st] OR “psychology”[st] OR “psychopharmacology”[st] OR “psychophysiology”[st] OR “public health”[st] OR “pulmonary medicine”[st] OR “radiology”[st] OR “radiotherapy”[st] OR “reproductive medicine”[st] OR “rheumatology”[st] OR “science”[st] OR “sexually transmitted diseases”[st] OR “social sciences”[st] OR “speech language pathology”[st] OR “sports medicine”[st] OR “statistics as topic”[st] OR “substance related disorders”[st] OR “technology”[st] OR “teratology”[st] OR “therapeutics”[st] OR “toxicology”[st] OR “transplantation”[st] OR “traumatology”[st] OR “tropical medicine”[st] OR “urology”[st] OR “vascular diseases”[st] OR “veterinary medicine”[st] OR “virology”[st] OR “vital statistics”[st] OR “women’s health”[st] OR “zoology”[st])

## Web-Appendix 9

Gini index calculation formula:

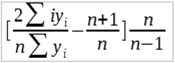

where n is the total number of authors, and yi is the number of journal articles published by author i, and authors are-sorted in non-decreasing order of journal article numbers.

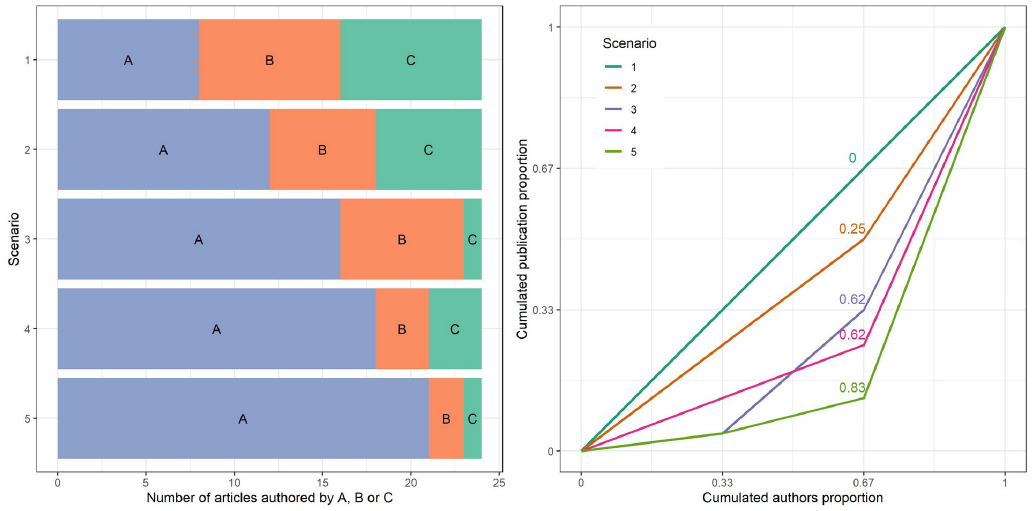

Figure: Theoretical cases of distribution of authorship between 3 authors totalizing 24 articles (left), with corresponding Lorenz curve and Gini index (right). The Lorenz curve is a visual representation of heterogeneity in the distribution of authorship, with a perfect equality producing a straight line between 0 and 1. The Gini index is the area between the line of equality and the Lorenz curve, corrected for the number of authors (n/n-1).

The Gini formula counts the times an author’s name occurs, but does not distinguish between papers contributed by different authors. Thus the first situation shown here could occur with 8 articles published in common by the 3 authors, or 4 articles published in common and each of the 3 separately publishing 4 other articles.

